# Connectomes, simultaneous EEG-fMRI resting-state data and brain simulation results from 50 healthy subjects

**DOI:** 10.1101/2024.04.17.589718

**Authors:** Jil Meier, Paul Triebkorn, Michael Schirner, Petra Ritter

## Abstract

We present raw and processed multimodal empirical data as well as simulation results from a study with The Virtual Brain (TVB).

Simultaneous electroencephalography (EEG) - functional magnetic resonance imaging (fMRI) resting-state data, diffusion-weighted MRI, and structural MRI were acquired for 50 healthy adult subjects (18 - 80 years of age) at the Charité University Medicine, Berlin, Germany.

We constructed personalized models from this multimodal data with TVB by optimizing parameters on an individual basis that predict multiple empirical features in fMRI and EEG, e.g. dynamic functional connectivity and bimodality in the alpha band power.

We annotated this large comprehensive empirical and simulated dataset according to the openMINDS metadata framework and structured it following Brain Imaging Data Structure (BIDS) standards for EEG and MRI as well as the BIDS Extension Proposal for computational modeling data.

This dataset provides ready-to-use data for future research at various levels of processing including the thereof inferred brain simulation results for a large dataset of healthy subjects with a wide age range.

## BACKGROUND AND SUMMARY

### Data origin

A description of the dataset and applied methods can be found in^1^. Please refer to this publication when re-using the simulated data. The acquisition and processing of the empirical data is described in our previous publications^2–4^, please cite these articles when using the presented raw and processed data.

Study participants joined voluntarily and experienced no cognitive, neurological or psychiatric conditions prior to this study based on self-reporting. All participants provided informed consent prior to entering the study. The research was performed in accordance with the Code of Ethics of the World Medical Association Declaration of Helsinki and after its approval by the local ethics committee at Charité University Berlin (application number EA1/041/13).

The presented dataset includes demographic (Table 1), imaging, electrophysiological as well as brain simulation data for N=50 subjects. More specifically, for each subject, we have diffusion-weighted magnetic resonance imaging (dwMRI), structural MRI, fieldmaps for distortion correction and simultaneous 22-minute resting-state electroencephalography (EEG)-functional MRI (fMRI) data as well as derivatives thereof. The derivatives are empirical structural and functional connectivity matrices, and BOLD time series aggregated according to the Desikan-Killiany parcellation and brain simulation data. More details can be found in Table 2.

**Table 1.**
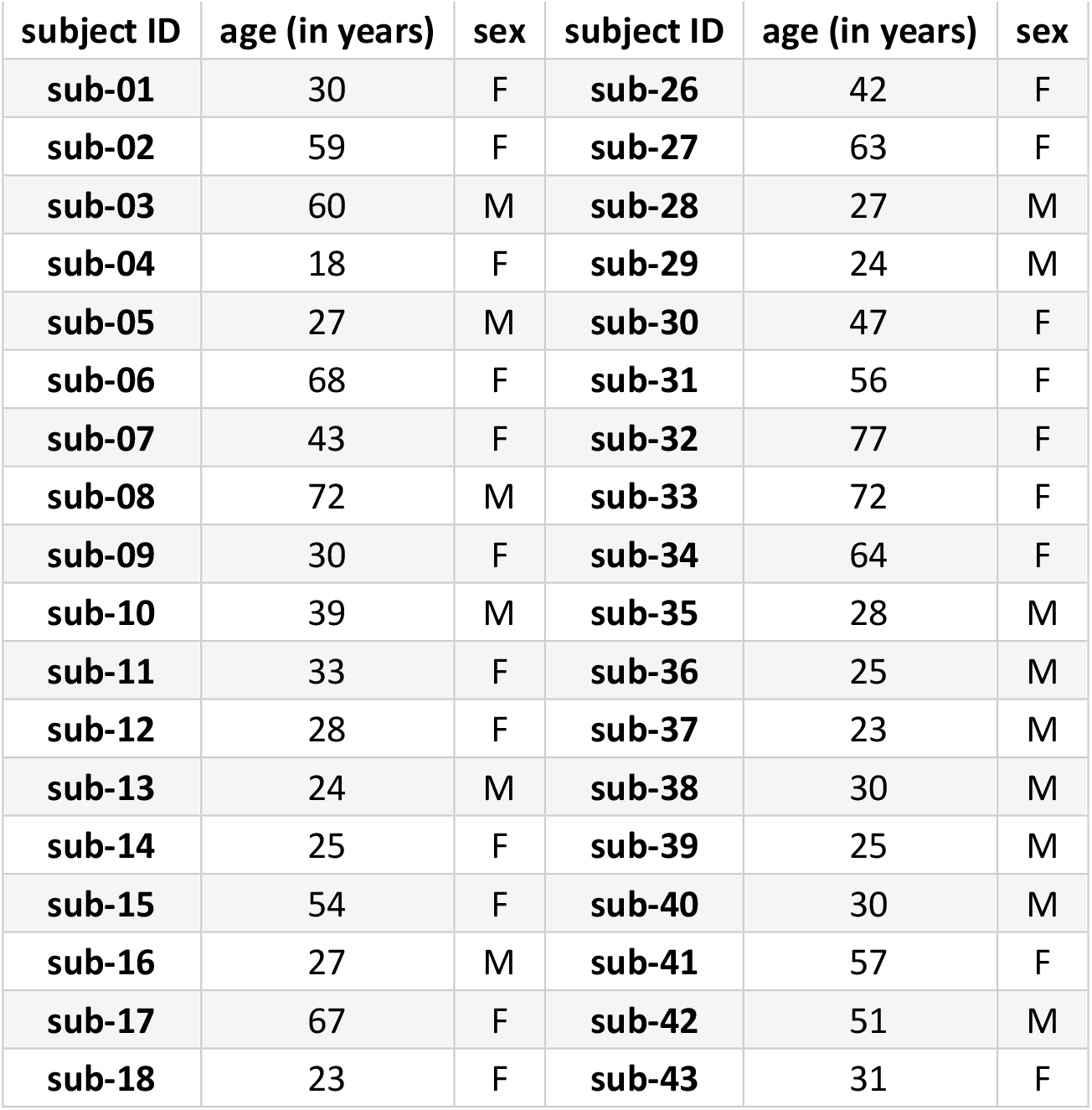
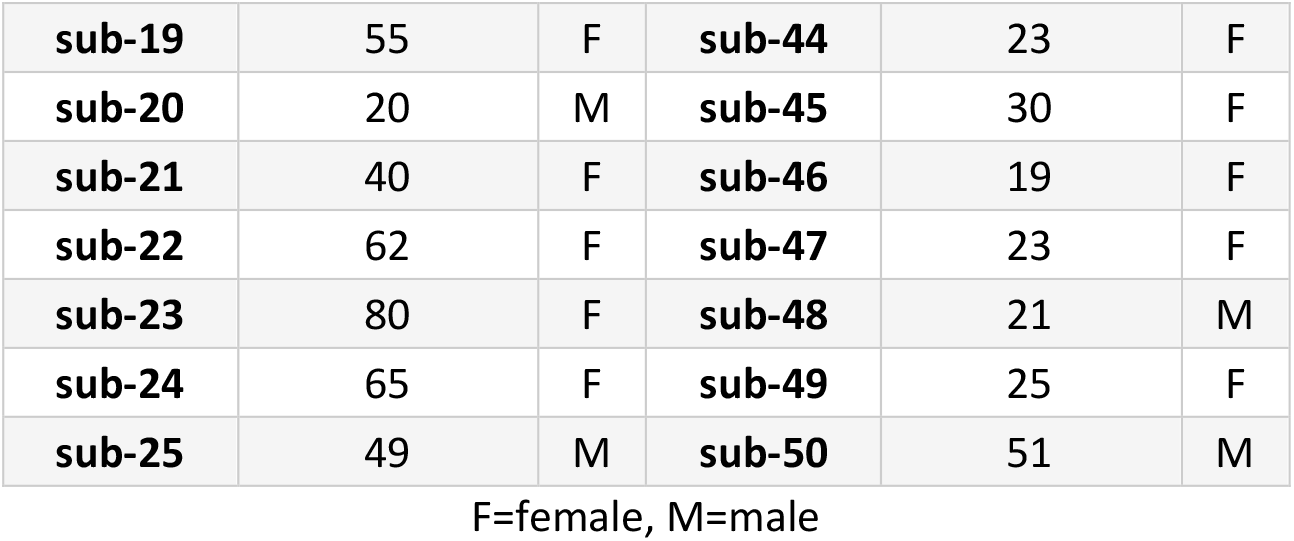
Subject overview. Subject identifier alongside age in years and sex are listed for all 50 subjects.

**Table 2.** Data overview. [number of subjects N for which this measurement data is available]

| **Data** | **Details** |
| --- | --- |
| Demographics | age [N=50], sex [N=50]: 18 - 80 years of age, mean 41.24±18.33; 31 females, 19 males; weight (for all subjects) and size (for 7 subjects) |
| **Modality** |  |
| **MRI sequences** |  |
| diffusion-weighted MR echo-planar measurements [N=50] | TR=7500ms, TE=86ms, FoV=220mm, 96 matrix, voxel size =2.3×2.3×2.3mm, no. of transversal slices=61 with thickness =2mm; 64 diffusion gradient directions with b-values =1000s/mm<sup>2</sup> |
| T1-weighted imaging [N=50] | MPRAGE sequence with 1x1x1mm T1-weighted imaging (TR = 1900ms, TE = 2.25ms, FA = 98, field of view (FoV) = 256mm, 256 matrix, no. of sagittal slices = 192 with thickness = 1mm) |
| EPI T2* [N=50] | 666 volumes, TR= 1940ms, TE=30ms, FA=788, FoV=192, 64 matrix, voxel size =3×3×3mm<sup>3</sup>, no. of transversal slices = 32 with thickness =3mm |
| functional MRI (fMRI) data [N=50] | 22min, resting state, simultaneous with EEG data |
| **EEG data** |  |
| electroencephalography (EEG) data [N=50] | 22 min, resting state, simultaneous with fMRI data |
| **Derivatives** |  |
| Structural connectivity [N=50] | distance and weight matrices (dimension: 68x68) and regional center coordinates for Desikan-Killiany atlas<sup>5</sup> |
| Empirical functional connectivity [N=50] | in Desikan-Killiany atlas parcellation<sup>5</sup>, based on empirical BOLD data; dimension: 68x68 |
| Empirical BOLD time series | in Desikan-Killiany atlas parcellation<sup>5</sup> |
| **Brain simulation data** |  |
| simulated BOLD and neural time series [N=50] | alpha parameter set: G=0.025:0.0001:0.04 and speed=10:10:100 simulated for 5 min; delta parameter set: G=0.05:0.01:0.25 and speed=20:20:100 simulated for 3 min |
| simulated BOLD FC [N=50] | Both for alpha and delta parameter set |

### Overview of previously published articles using this data

The data was acquired in the lab of Petra Ritter at Charité University Medicine Berlin. As the earliest publication using this dataset, Ritter et al.^2^ introduced the used brain modeling software, The Virtual Brain (TVB, thevirtualbrain.org), and used an initial subset of the presented data, namely nine subjects (mean age 24.6 years, five men), to provide a proof-of-principle for TVB. A different parcellation was used, the monkey brain surface^6^ brought into MNI space^7,8^. All subjects were fitted with the Stefanescu-Jirsa three-dimensional model (SJ3D) model to their empirical EEG and blood oxygen level dependent (BOLD) signal data.

Schirner et al.^3^ presented the used processing pipeline and applied it to 49 subjects (age range 18–80 years, 30 females, 19 males). Zimmermann et al.^9^ analyzed structural connectivity (SC) - functional connectivity (FC) couplings of 47 subjects (age range 18-80, mean age ± SD= 41.49 ± 18.36; 19 male/28 female) using their BOLD data and individual connectomes. They found an age effect of decreasing FC and SC in most brain areas and SC-FC coupling on the level of regions predicts age superior to single FC or SC. Deco and coauthors^10^ constructed individual whole-brain models by fitting FC and functional connectivity dynamics (FCD) out of 24 subjects (age range 18 – 33 years, mean age 25.7, 12 females, 12 males) using a normal form of a supercritical Hopf bifurcation and showed that the models recreate resting-state activity best when exhibiting maximum metastability and revealed a dynamical core that the activity of the whole brain. The authors of Glomb et al.^11^ analyzed the FCD of 24 subjects (age range 18-35 years, mean age = 25.7 years, 11 females, 13 males) by applying tensor decomposition and identified communities that resemble resting-state networks. These resting-state networks also emerge from simulated data with the reduced Wong-Wang model^12^. Schirner et al.^4^ used a subset of 15 subjects (age range: 18–31 years, 8 females) for their analysis. They selected the youngest 15 subjects from the larger dataset of 49 subjects used in^3^ to ensure highest quality for EEG recordings after applying the MR artefact corrections. All subjects were fitted with the reduced Wong-Wang model^12,13^ to their individual regional time series based on BOLD fMRI. The fitted model was then driven with individual empirical EEG source activity instead of noise and yielded various known empirical phenomena from literature. In Zimmermann et al.^14^, using a subset of 48 subjects (age range 18-80 years, mean age ± standard deviation (SD) = 41.9 ± 18.47), the authors showed that there is a correlation between group-averaged SC and FC but at the individual level all subject’s SCs correlated significantly with all FCs. Further an age effect in the FC but not in the SC was shown. The authors in Battaglia et al.^15^ analyzed the dynamic FC in the subset of N=49 subjects and demonstrated that it becomes more reandom and less complex with aging. On the same subset, another study constructed virtual connectomes based on whole-brain network modeling using stochastic linear models and reduced Wong-Wang models^13,16^ and showed their suitability for replacing empirical data in distinguishing age classes^17^. Moreover, Goldman et al.^18^ used the subject sub-09 for the underlying connectome, simulated the Adaptive-Exponential Integrate-and-Fire (AdEx) model for wakefulness and slow-wave sleep and performed state-dependent virtual transcranial magnetic stimulation (TMS). In a follow-up study, Goldman et al.^19^ used the subject sub-36 to test the AdEx mean-field model on its connectome. Tesler et al.^20^ used the data of the sub-36 to test a newly proposed method of local field potential (LFP) and magnetoencephalography (MEG) signal forward modeling based on mean-field models. They also used the AdEx model as in^19^.

### Data use and sharing conditions

Use and access of the dataset is governed by the permissions, rules and obligations set out in the accompanying

- Data Processing Agreement (DPA, please see below)
- Creative Commons License

We provide a pre-filled DPA for the dataset (EU Standard Contractual Clauses, https://commission.europa.eu/law/law-topic/data-protection/international-dimension-data-protection/standard-contractual-clauses-scc_en, https://wiki.ebrains.eu/bin/view/Collabs/simultaneous-eeg-fmri/).

The agreement is made between controllers and processors of the dataset, mutually assuring compliance with the General Data Protection Regulation.

The data is shared via the Virtual Research Environment (VRE, https://vre.charite.de) at Charité – Universitätsmedizin Berlin, a node of EBRAINS Health Data Cloud. Prospective processors fill the pre-filled DPA as described below and send it to the controller to form a data processing agreement as legal basis for processing under GDPR.

Prospective Processors fill the pre-filled DPA in the Drive of this Collab with their information. Specifically, they enter

○ Annex I:
  ▪ Names and Addresses of all Prospective Processors
  ▪ Contact information of the responsible institutional Data Protection Officer
  ▪ Signature (either by printing and scanning or with a valid digital signature)
○ Annex II: Duration of the processing
○ Annex III: Prospective Processors add to the list of technical and organizational measures carried out to ensure the security of the data, particularly regarding the safety of the processing after downloading and decrypting the data.

Prospective Processors send the signed document to the controller which will evaluate and sign the agreement and arrange the transmission of the data.

We explicitly grant re-use of the overall dataset under the Creative Commons License Attribution-ShareAlike 4.0 International. The license explicitly grants reuse of all non-personal aspects of the dataset. Importantly, the personal data contained within this dataset is governed by the EU General Data Protection Regulation. As a consequence, the license only applies to non-personal aspects of the dataset, for example, relating to the structure and organization of the dataset or the way the dataset was produced, but not to the personal information contained within the dataset.

Data and code sharing will be possible only after journal acceptance.

## METHODS

These methods are expanded versions of descriptions in our related works^1–4^.

### Data acquisition

Scanning was performed on a Siemens Tim Trio MR scanner (12-channel Siemens head coil). First, anatomical and dwMRI sequences were run and then participants had their EEG cap set up outside of the scanner. Subsequently, the simultaneous EEG-fMRI measurements were performed with a total length of 22 min. Instructions during this time were to close their eyes, relax but not fall asleep, which are denoted as resting-state scans.

An MPRAGE sequence with 1×1×1mm T1-weighted imaging was performed (repetition time (TR) = 1900ms, echo time (TE) = 2.25ms, flip angle (FA) = 98, field of view (FoV) = 256mm, 256 matrix, no. of sagittal slices = 192 with thickness = 1mm). Moreover, an echo planar imaging (EPI) T2* sequence was run (666 volumes, TR= 1940ms, TE=30ms, FA=788, FoV=192, 64 matrix, voxel size =3×3×3mm^3^, no. of transversal slices = 32 with thickness =3mm). The details of the diffusion-weighted MR echo-planar measurements were TR=7500ms, TE=86ms, FoV=220mm, 96 matrix, voxel size =2.3×2.3×2.3mm, no. of transversal slices=61 with thickness =2mm; 64 diffusion gradient directions with b-values =1000s/mm^2^. We removed the first five images of the BOLD fMRI scan due to saturation effects.

### Preprocessing

Our pipeline for preprocessing connectomes^3^ was used. For the T1-weighted images, we performed motion correction, intensity normalization, extraction of non-brain tissue, brain mask generation, and segmentation of cortical and subcortical grey matter. The Desikan-Killiany Atlas^5^ from FREESURFER was used as parcellation resulting in 68 regions. In addition, we carried out a manual quality check of the parcellation for individual high-resolution T1-weighted scans.

As steps to preprocess the dwMRI data, we implemented motion and eddy current correction as well as linearly registered the b0 image to the individual T1-weighted image. The parcellation was brought to individual diffusion space before probabilistic tractography was applied. To constrain tractography, spherical deconvolution was conducted with MRTrix streaming method being able to identify crossing fibers (fractional anisotropy threshold =0.1)^21^. We meticulously sampled the gray matter – white matter interface and streamlined up to 200,000 times from each voxel (radius of curvature =1mm, maximum length =300mm). As a result of the tractography, the SC (68×68) matrix was computed for each region pair. The region labels can be found in Table 3. All subjects have the complete file collection listed in the Data Records section below.

**Table 3.** Region labels. Labels for the 68 regions, using the same order as in the connectome. The prefixes “lh_…” and “rh_…” specify left and right hemisphere, respectively. Table taken from^1^ (Supplementary Table 1 there).

| Number | Region label | Number | Region label |
| --- | --- | --- | --- |
| 1 | lh_bankssts | 35 | rh_bankssts |
| 2 | lh_caudalanteriorcingulate | 36 | rh_caudalanteriorcingulate |
| 3 | lh_caudalmiddlefrontal | 37 | rh_caudalmiddlefrontal |
| 4 | lh_cuneus | 38 | rh_cuneus |
| 5 | lh_entorhinal | 39 | rh_entorhinal |
| 6 | lh_fusiform | 40 | rh_fusiform |
| 7 | lh_inferiorparietal | 41 | rh_inferiorparietal |
| 8 | lh_inferiortemporal | 42 | rh_inferiortemporal |
| 9 | lh_isthmuscingulate | 43 | rh_isthmuscingulate |
| 10 | lh_lateraloccipital | 44 | rh_lateraloccipital |
| 11 | lh_lateralorbitofrontal | 45 | rh_lateralorbitofrontal |
| 12 | lh_lingual | 46 | rh_lingual |
| 13 | lh_medialorbitofrontal | 47 | rh_medialorbitofrontal |
| 14 | lh_middletemporal | 48 | rh_middletemporal |
| 15 | lh parahippocampal | 49 | rh parahippocampal |
| 16 | lh_paracentral | 50 | rh_paracentral |
| 17 | lh_parsopercularis | 51 | rh_parsopercularis |
| 18 | lh_parsorbitalis | 52 | rh_parsorbitalis |
| 19 | lh_parstriangularis | 53 | rh_parstriangularis |
| 20 | lh_pericalcarine | 54 | rh_pericalcarine |
| 21 | lh_postcentral | 55 | rh_postcentral |
| 22 | lh_posteriorcingulate | 56 | rh_posteriorcingulate |
| 23 | lh_precentral | 57 | rh_precentral |
| 24 | lh_precuneus | 58 | rh_precuneus |
| 25 | lh_rostralanteriorcingulate | 59 | rh_rostralanteriorcingulate |
| 26 | lh_rostralmiddlefrontal | 60 | rh_rostralmiddlefrontal |
| 27 | lh_superiorfrontal | 61 | rh_superiorfrontal |
| 28 | lh_superiorparietal | 62 | rh_superiorparietal |
| 29 | lh_superiortemporal | 63 | rh_superiortemporal |
| 30 | lh_supramarginal | 64 | rh_supramarginal |
| 31 | lh_frontalpole | 65 | rh_frontalpole |
| 32 | lh_temporalpole | 66 | rh_temporalpole |
| 33 | lh_transversetemporal | 67 | rh_transversetemporal |
| 34 | lh_insula | 68 | rh_insula |

**Table 4.** List of all file formats. Together with an explanation which software created the respective files.

| Format | Extension | Software used / file specification |
| --- | --- | --- |
| Tab-Separated Value | tsv | Generated by semi-customized Matlab script `/comp/code/data_compmodel_into_BIDS.m` for `/comp` files, by bidscoin (https://github.com/Donders-Institute/bidscoin)<sup>32</sup> for `/mri` files and by MNE-BIDS<sup>33</sup> for `/eeg` files |
| JavaScript Object Notation | json | Generated by semi-customized Python script `/comp/code/create_jsons_for_BIDS-EP_comp_model_data.ipynb` for `/comp` files, by bidscoin (https://github.com/Donders-Institute/bidscoin)<sup>32</sup> for `/mri` files and by MNE-BIDS<sup>33</sup> for `/eeg` files |
| NIfTI file | nii.gz | bidscoin (https://github.com/Donders-Institute/bidscoin)<sup>32</sup> |
| B value file | bval | raw data from MRI scanner |
| B vector file | bvec | raw data from MRI scanner |
| Raw EEG data file | eeg | BrainVision Recorder software |
| Header EEG file | vhdr | BrainVision Recorder software |
| Marker EEG file | vmrk | BrainVision Recorder software |
| Extensible Markup Language | xml | Self-made in LEMS format generated by semi-customized Python script `/comp/code/create_jsons_for_BIDS-EP_comp_model_data.ipynb` |

If a region pair had at least one track between them, then the number of voxel pairs between those two regions was denoted as their link weight and put into the weight matrix. The weight matrix was further transformed by taking the common logarithm and normalizing the entries between 0 and 1, i.e. dividing by its maximum value. The distance matrix represents the measured average track length of the fibers (in mm) between all region pairs with respect to their centers.

One of our previous articles explains the details of preprocessing simultaneous EEG-fMRI data^22^. The BOLD data went through the following preprocessing steps: motion correction, brain extraction, high-pass filter (100s), registration to individual T1-weighted scans, application of high-resolution parcellation mask. Temporal signal-to-noise maps^23^ were used to monitor BOLD signal quality. For each region, we averaged across voxel to obtain the regional BOLD signal. The functional connectivity matrix consists of pairwise Pearson’s correlation coefficients between BOLD data.

The EEG recording system was an MR-compatible 64-channel system (BrainAmp MR Plus; Brain Products) together with an MR-compatible EEG cap (Easy Cap) including ring-type sintered silver chloride electrodes with iron-free copper leads. We arranged 61 scalp electrodes following the International 10-20 System with impedance of all electrodes < 15kΩ and each electrode had an impedance of 10kΩ to avoid heating due to magnetic field switches. Reference electrode was FCz and we recorded also two electrocardiograms and one electrooculography (EOG) channel. EEG amplifier recorded at a range of ±16.38 MV, resolution 0.5 μV and 5 kHz sampling rate. In addition, a hardware embedded low-pass filter of 250 Hz was applied. A synchronization between EEG sampling clock and gradient-switching clock of the MR scanner was performed prior to recording^24^. EEG preprocessing included image-acquisition artifact and ballistrocardiogram correction and was described in^3,4,24–29^.

### Brain Simulation

All computational modeling data can be re-generated by following the processing steps described in Methods and in^1^. Inside TVB, the neural mass model of Stefanescu-Jirsa 3D was chosen to represent regional activity. Simulations were run for generating both the alpha and the delta frequency rhythm with different local parameters, alpha and delta parameter set, respectively. Brain regions were coupled with global coupling factor G and conduction speed v. Depending on the alpha or delta parameter set, different ranges of G and v were tested and simulated (alpha parameter set: G=0.025:0.0001:0.04 and speed=10:10:100, thus 150×10=1,500 parameter combinations, simulated for 5 min; delta parameter set: G=0.05:0.01:0.25 and speed=20:20:100, thus 20×5=100 parameter combinations, simulated for 3 min). These simulations were resulting in two types of generated time series, the simulated BOLD and neural time series. Outside TVB, the regional BOLD time series were analyzed with respect to their pairwise correlations and a simulated FC matrix was obtained. This simulated FC matrix was compared with the empirical counterpart, which was generated based on empirical regional BOLD time series.

Optimal global parameters of G and v were chosen based on the optimal fit between empirical and simulated FC. This data flow process was the same for all N=50 subjects. All statistical tests used for the analysis of empirical and simulated data are listed in the previous publication^1^.

### Data standardization into BIDS format

We transformed all different kinds of collected and generated data into the Brain Imaging Data Structure (BIDS) format^30^. Thereby, we followed the BIDS standard for MRI data^30^, for functional MRI derivatives (BIDS extension proposal (BEP) 012, https://github.com/bids-standard/bids-specification/pull/519) to convert the empirical BOLD time series, for EEG data^31^, and for computational modeling data recently proposed by (Schirner & Ritter, 2023, https://zenodo.org/record/7962032). For the standardization of empirical data, we used existing BIDS converters. The DICOM MRI data was transformed to *NIfTI* format and saved according to BIDS specifications using bidscoin (https://github.com/Donders-Institute/bidscoin)^32^. In the prepared data records, we also share the code in the form of the corresponding *.yaml* file that was used for conversion, which will be made public after publication.

For the conversion of EEG data to BIDS, we used the package MNE-BIDS^33^. All of the empirical BIDS-formatted data was validated with the BIDS validator (https://github.com/bids-standard/bids-validator) (Blair et al., 2022, https://zenodo.org/record/6391626). The used code is shared in the code subfolder of the EEG data folder in the form of a Jupyter notebook and will be published after journal acceptance.

We wrote semi-customized scripts in Matlab and Python for conversions of the computational modeling data into the BIDS format as suggested by the BEP for computational modeling data (Schirner & Ritter, 2023, https://zenodo.org/record/7962032), which are also shared in the code folder of our data structure (and will be published after journal acceptance). The conversion involved creating the folder structure, creating the *json* sidecar files and filling them with all available metadata, the conversion of existing *.mat* files into *.tsv* files, renaming of all files and folders and compression of *.tsv* files. In addition, we created an *.xml* file in LEMS format for the SJ3D model. We are currently developing an app called *sim2bids* (https://github.com/BrainModes/sim2bids/) for automatic conversion of computational modeling data to BIDS, generalizing these conversion scripts in a convenient GUI format to multiple data storage formats (.h5 files, .mat files, .pkl files etc.).

For the openMINDS metadata annotation, we used the EBRAINS Metadata Wizard (https://metadata-wizard.apps.ebrains.eu/) and made the data available in the EBRAINS Knowledge Graph in three different formats (derived (Meier et al., 2025a, https://doi.org/10.25493/6CKF-MJS), simulated (Meier et al., 2025b, https://doi.org/10.25493/R7DJ-3NQ) and raw data (Meier et al., 2025c, https://doi.org/10.25493/RSFP-PS6).

## DATA RECORDS

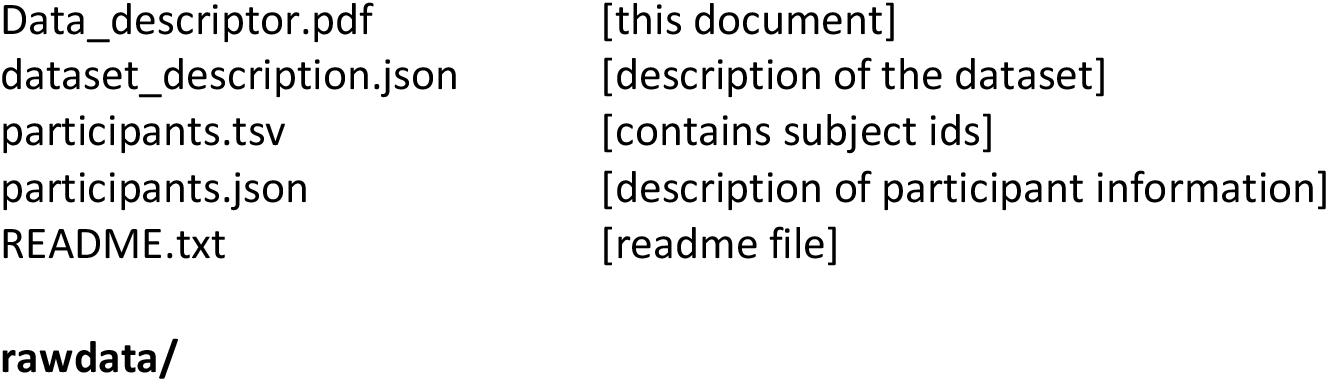
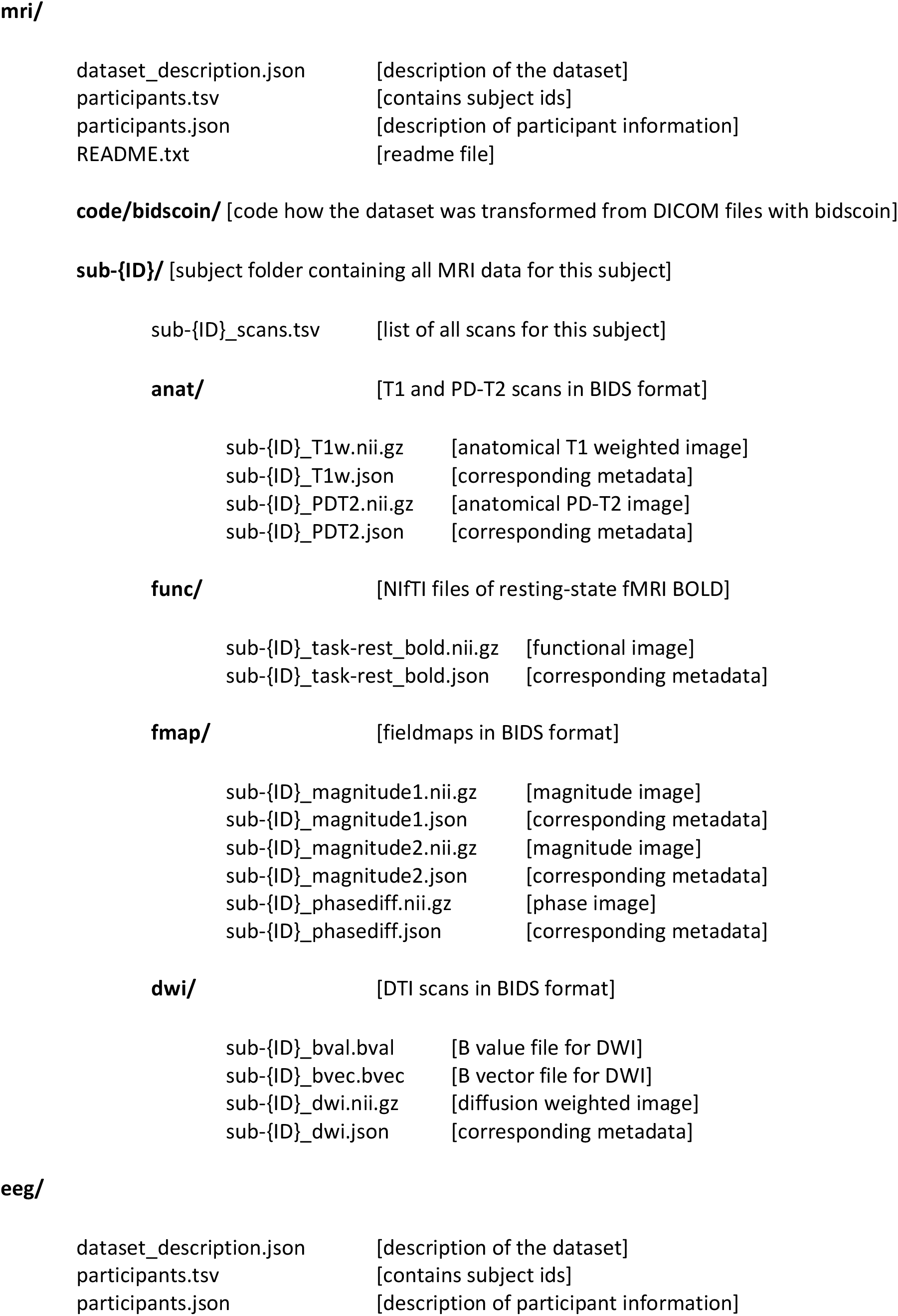
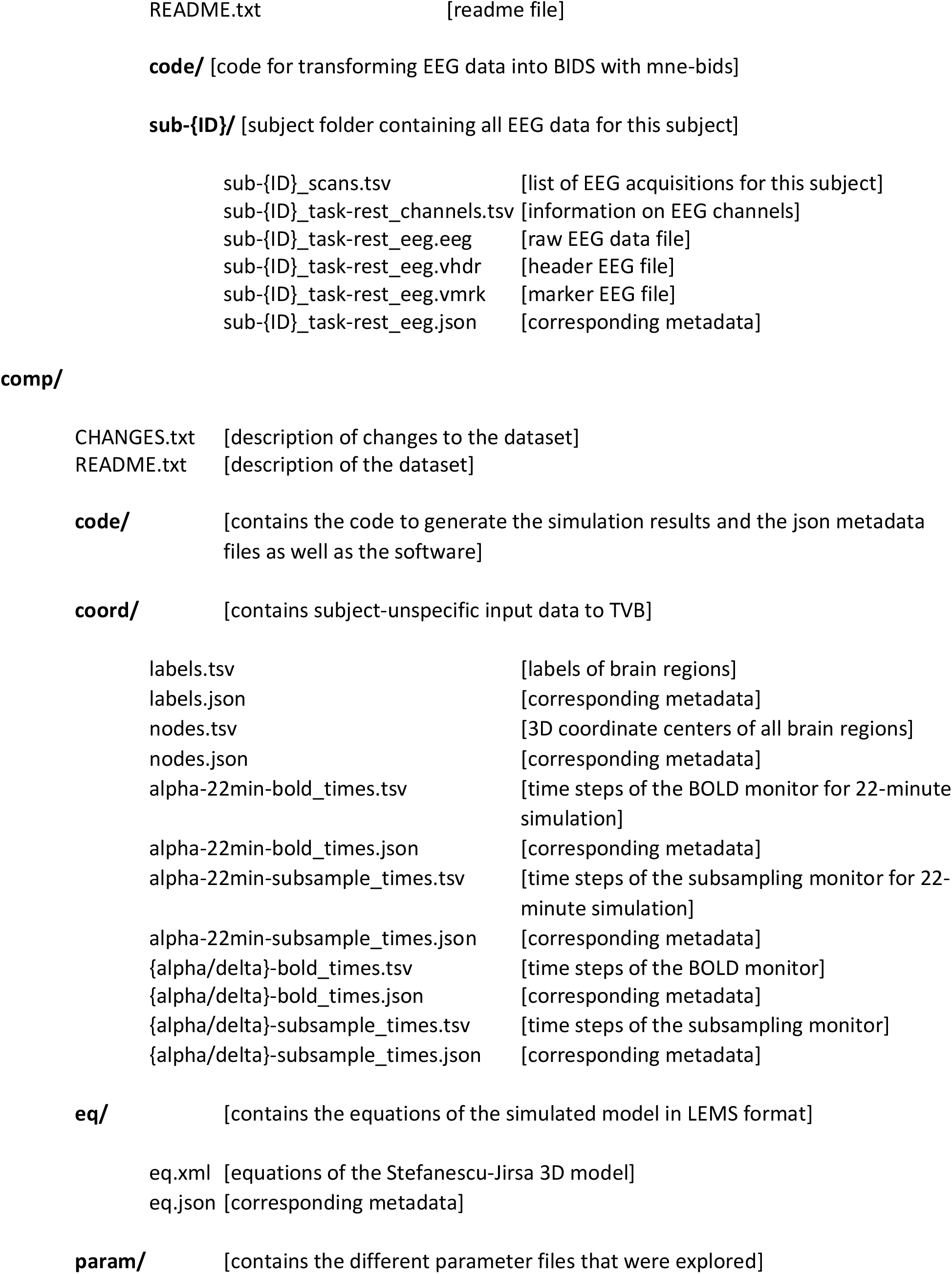
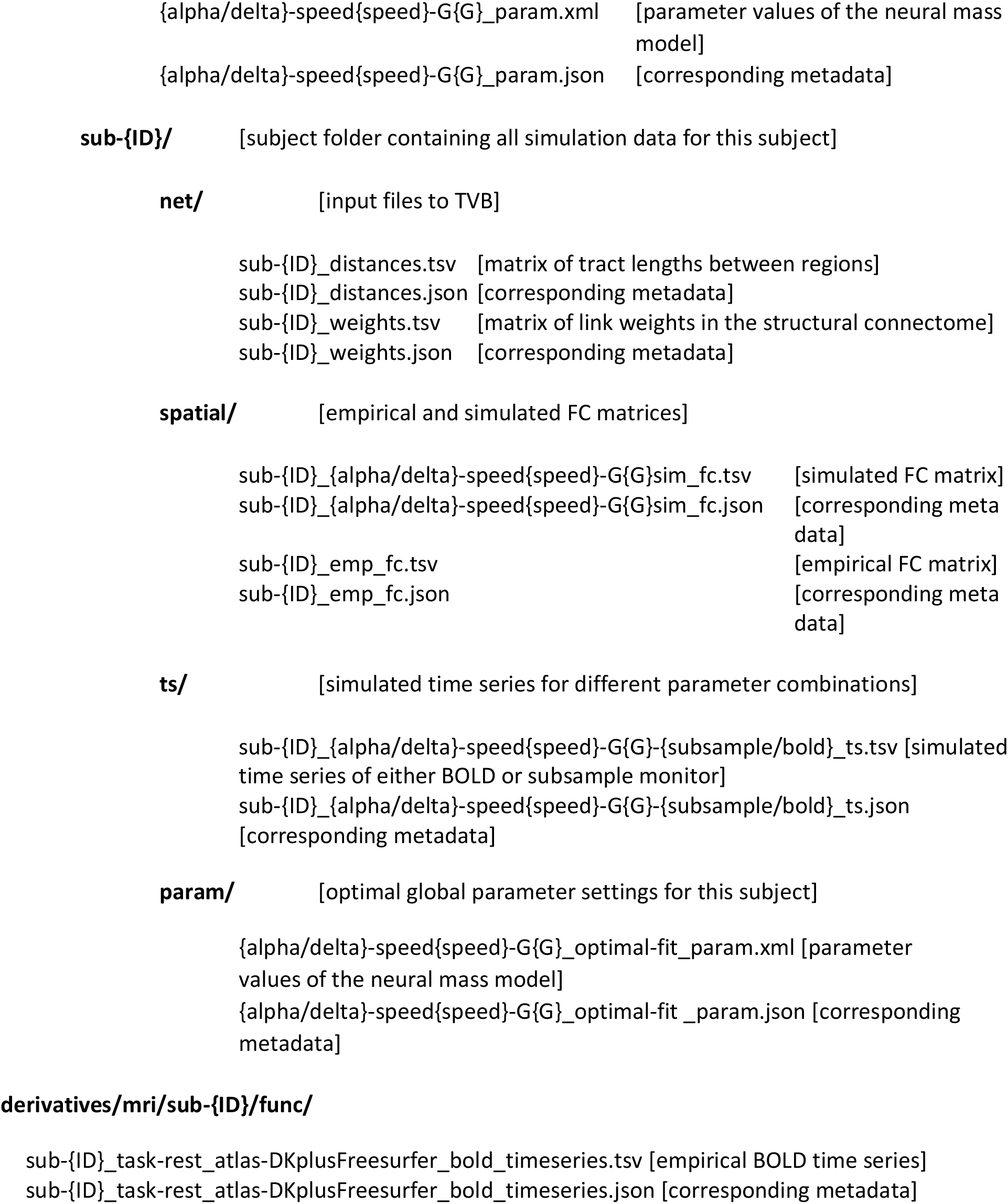

|  |  |
| --- | --- |
| Data_descriptor.pdf | [this document] |
| dataset_description.json | [description of the dataset] |
| participants.tsv | [contains subject ids] |
| participants.json | [description of participant information] |
| README.txt | [readme file] |
| dataset_description.json | [description of the dataset] |
| participants.tsv | [contains subject ids] |
| participants.json | [description of participant information] |
| README.txt | [readme file] |
| sub-{ID}_T1w.nii.gz | [anatomical T1 weighted image] |
| sub-{ID}_T1w.json | [corresponding metadata] |
| sub-{ID}_PDT2.nii.gz | [anatomical PD-T2 image] |
| sub-{ID}_PDT2.json | [corresponding metadata] |
| sub-{ID}_task-rest_bold.nii.gz | [functional image] |
| sub-{ID}_task-rest_bold.json | [corresponding metadata] |
| sub-{ID}_magnitude1.nii.gz | [magnitude image] |
| sub-{ID}_magnitude1.json | [corresponding metadata] |
| sub-{ID}_magnitude2.nii.gz | [magnitude image] |
| sub-{ID}_magnitude2.json | [corresponding metadata] |
| sub-{ID}_phasediff.nii.gz | [phase image] |
| sub-{ID}_phasediff.json | [corresponding metadata] |
| sub-{ID}_bval.bval | [B value file for DWI] |
| sub-{ID}_bvec.bvec | [B vector file for DWI] |
| sub-{ID}_dwi.nii.gz | [diffusion weighted image] |
| sub-{ID}_dwi.json | [corresponding metadata] |
| **eeg/** |  |
| dataset_description.json | [description of the dataset] |
| participants.tsv | [contains subject ids] |
| participants.json | [description of participant information] |
| sub-{ID}_scans.tsv | [list of EEG acquisitions for this subject] |
| sub-{ID}_task-rest_channels.tsv | [information on EEG channels] |
| sub-{ID}_task-rest_eeg.eeg | [raw EEG data file] |
| sub-{ID}_task-rest_eeg.vhdr | [header EEG file] |
| sub-{ID}_task-rest_eeg.vmrk | [marker EEG file] |
| sub-{ID}_task-rest_eeg.json | [corresponding metadata] |
| labels.tsv | [labels of brain regions] |
| labels.json | [corresponding metadata] |
| nodes.tsv | [3D coordinate centers of all brain regions] |
| nodes.json | [corresponding metadata] |
| alpha-22min-bold_times.tsv | [time steps of the BOLD monitor for 22-minute simulation] |
| alpha-22min-bold_times.json | [corresponding metadata] |
| alpha-22min-subsample_times.tsv | [time steps of the subsampling monitor for 22-minute simulation] |
| alpha-22min-subsample_times.json | [corresponding metadata] |
| {alpha/delta}-bold_times.tsv | [time steps of the BOLD monitor] |
| {alpha/delta}-bold_times.json | [corresponding metadata] |
| {alpha/delta}-subsample_times.tsv | [time steps of the subsampling monitor] |
| {alpha/delta}-subsample_times.json | [corresponding metadata] |
| {alpha/delta}-speed{speed}-G{G}_param.xml | [parameter values of the neural mass model] |
| {alpha/delta}-speed{speed}-G{G}_param.json | [corresponding metadata] |
| sub-{ID}_distances.tsv | [matrix of tract lengths between regions] |
| sub-{ID}_distances.json | [corresponding metadata] |
| sub-{ID}_weights.tsv | [matrix of link weights in the structural connectome] |
| sub-{ID}_weights.json | [corresponding metadata] |
| sub-{ID}_{alpha/delta}-speed{speed}-G{G}sim_fc.tsv | [simulated FC matrix] |
| sub-{ID}_{alpha/delta}-speed{speed}-G{G}sim_fc.json | [corresponding meta data] |
| sub-{ID}_emp_fc.tsv | [empirical FC matrix] |
| sub-{ID}_emp_fc.json | [corresponding meta data] |
| sub-{ID}_{alpha/delta}-speed{speed}-G{G}-{subsample/bold}_ts.tsv | [simulated time series of either BOLD or subsample monitor] |
| sub-{ID}_{alpha/delta}-speed{speed}-G{G}-{subsample/bold}_ts.json | [corresponding metadata] |
| {alpha/delta}-speed{speed}-G{G}_optimal-fit_param.xml | [parameter values of the neural mass model] |
| {alpha/delta}-speed{speed}-G{G}_optimal-fit_param.json | [corresponding metadata] |
| **derivatives/mri/sub-{ID}/func/** |  |
| sub-{ID}_task-rest_atlas-DKplusFreesurfer_bold_timeseries.tsv | [empirical BOLD time series] |
| sub-{ID}_task-rest_atlas-DKplusFreesurfer_bold_timeseries.json | [corresponding metadata] |

Additionally, files can be labelled with “acquisition” or “run” label for multiple files of MRI or EEG data. Complemenary to these data records, we generated an interactive *html* file, providing a preview of all of the prepared filenames in an interactive file tree (Supplementary Information). Moreover, we prepared the data of one example subject (sub-01) to be publicly shared (after acceptance) by deleting all identifiable files and selecting only a subset of simulated metadata files.

## TECHNICAL VALIDATION

Technical validation for this dataset was presented in a previous study^3^. We here present a short summary of these results.

It is important to have a robust pipeline for generating the individual connectomes, therefore Schirner et al.^3^ tested for intra- and inter-subject variability. For this purpose, three of the fifty subjects were scanned three times with their anatomical and diffusion weighted scans. The first two scans were acquired directly following each other (no break) and the third scan was after being shortly moved outside the scanner to modify the subject’s head position. Figure 1 displays boxplots of correlation coefficients when comparing strength and distance matrices among and within the three subjects. All of the different computations of the weight matrices had high correlation coefficients (raw counts (0.97– 0.99, 0.98 ± 0.007), distinct connections (0.97–0.99, 0.98 ± 0.006), and weighted distinct connections (0.96–0.98, 0.98 ± 0.007)). The similarity between subjects was significantly lower (raw counts (0.87– 0.93, 0.9 ± 0.02), distinct connections (0.87–0.92, 0.89 ± 0.02), and weighted distinct connections (0.86– 0.91, 0.89 ± 0. 01)) than the one within subjects. Matrices filled with distances exhibited lower similarities (0.84–0.92, 0.88 ± 0.03) than strength matrices but still had a higher intra-than inter-subject similarity (0.68–0.77, 0.72 ± 0.03).

**Figure 1.**
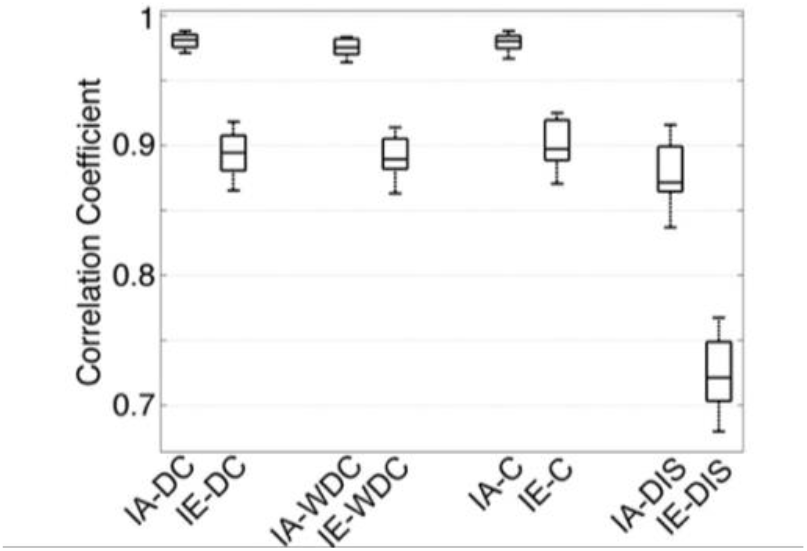
Intra-compared with inter-subject variability. Characterized by boxplots of all correlation coefficients (CC) between all pairs of strengths matrices of three subjects and three scans. Boxes labeled with ‘IA’ denote intra-subject CC, while ‘IE’ denotes inter-subject CC. DC = distinct connections, WDC = weighted distinct connections, C = raw counts, and DIS = distance. Figure adapted from Figure 4 in^3^.

In addition to correlations, the intra-class-coefficients (ICC)^34,35^ were analyzed for the three datasets from the three subjects to quantify the consistency among multiple scans (Table 5). More specifically, ICC(3,1) = (MC_R_ – MS_E_)/(MS_R_ + (k-1)MS_E_) was used, where MC_R_ is the mean square for rows of observables (i.e. strengths/distances between nodes and node degrees), MS_E_ the mean squared error and 1 denotes a perfect similarity between scans. The variable *k* denotes the number of observables. The ICC(3,1) was computed for all connectivity matrices and for node strengths, which is defined as the sum of all link weights connected to a node. Table 5 displays that we have an almost perfect agreement among the three scans within the same subjects. In the same line of analysis, a different version of ICC was used to determine whether we have a higher variability between than within subjects, ICC = σ^2^_bs_ / (σ^2^_bs_ + σ^2^_ws_), where σ^2^_bs_ describes the variance between subjects and σ^2^_ws_ the pooled one within the same subject. If this version of the ICC yields a higher ration than 0.5, it indicates that there is more variability between subjects than within the same subject. For each node, we computed the ICC providing an average ICC of for distinct connections of 0.77±0.16, for raw counts 0.8±0.13 and weighted distinct connections 0.76±0.15. These high values confirm that there is more variance between subjects than within the same subject. Moreover, the coefficient of variation (CV), which is defined as σ^2^_ws_ divided by the overall measurement mean^36^, was calculated to analyze the variability between scans and subjects of node strengths. If CV < 1, the variability is determined as low, for CV > 1 it is considered as high. In our case, the CV was low for node strengths, 0.07 ± 0.05 (distinct connections), 0.06 ± 0.03 (weighted distinct connections) and 0.05 ± 0.03 (raw counts), indicating robust computation of node strengths (Table 6).

**Table 5.** Test–retest analysis for pipeline generated SC matrices. Table taken from^3^.

| Subject | RC | DC | WDC | DIS | NS(RC) | NS(DC) | NS(WDC) | NS(DIS) |
| --- | --- | --- | --- | --- | --- | --- | --- | --- |
| 1 | 0.98 | 0.98 | 0.97 | 0.88 | 1 | 1 | 0.99 | 0.98 |
| 2 | 0.98 | 0.98 | 0.97 | 0.86 | 1 | 0.99 | 0.99 | 0.97 |
| 3 | 0.98 | 0.98 | 0.98 | 0.89 | 1 | 0.99 | 0.99 | 0.98 |

*Summary of test–retest analysis for the three capacities metric and the distance estimation. ICC(3,1) values were computed over full networks with matrices being thresholded at fixed values (thresholds: 4000 for DC, 4500 for RC, 100 for WDC). RC = raw counts, DC = distinct connections, WDC = weighted distinct connections, DIS = distances, and NS = node strength.*

**Table 6.**
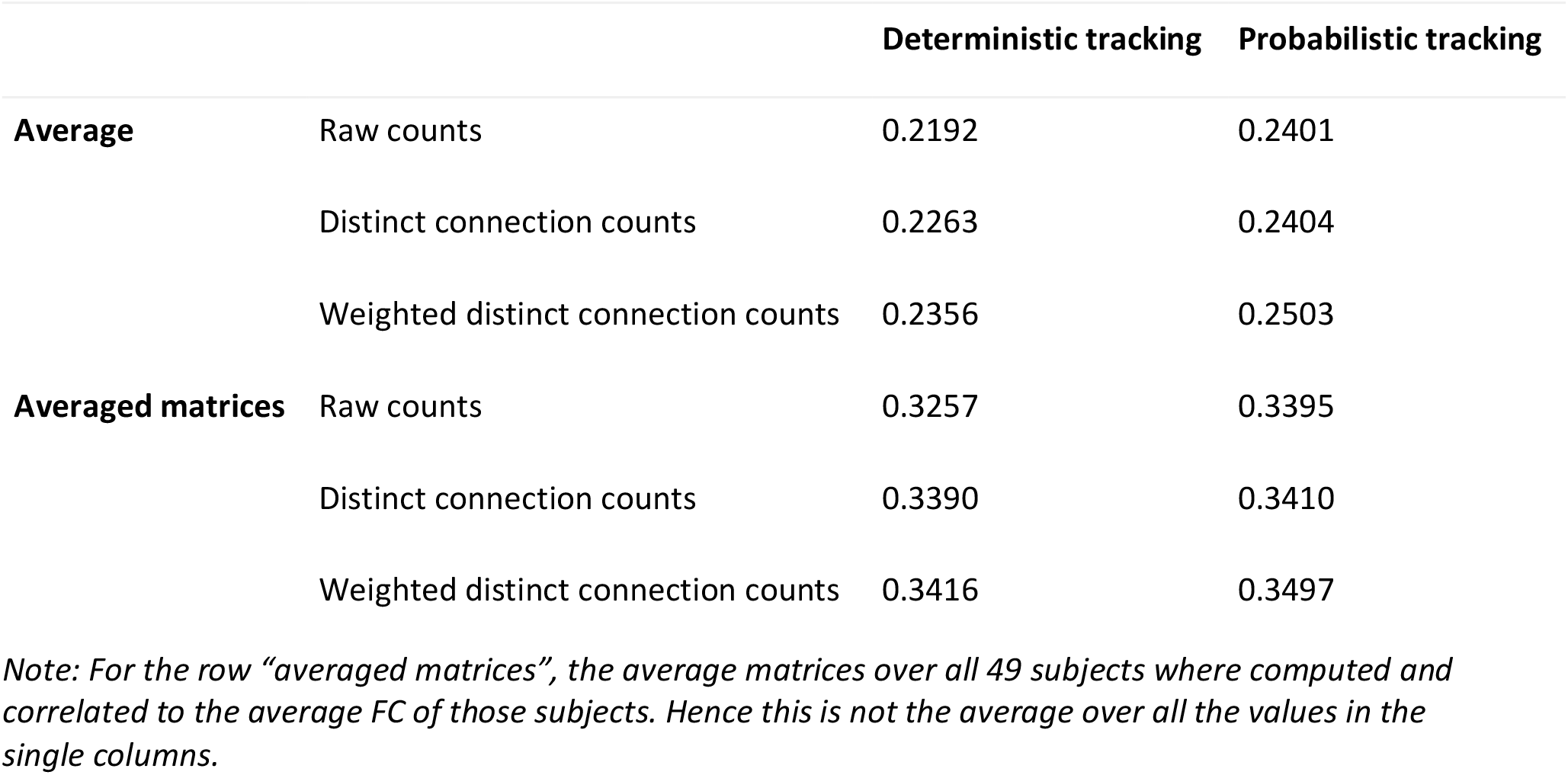
Correlation between SC and FC using different tracking methods. Table taken from^3^.

To sum up, for this small sample of three subjects, the test-retest reliability of structural connectome estimation was very high and in general, the intra-subject variability was smaller than the inter-subject variability. Compared to previous literature, the reproducibility of our connectomes was higher or similar (as in e.g.^37^). While we reported an average intra-subject correlation over all weighting schemes of r=0.98 and an ICC(3,1)>0.97, other studies obtained similar results: r=0.78 in^38^, r=0.89 and r=0.84 for different weight calculations in^39^, ICC(3,1) = 0.76 for global network strength and ICC(3,1)=0.62 for node strength in^40^.

As both SC and FC matrices are computed for each individual subject, we compared both modalities for the group-averaged matrices for different weight computations and different tracking algorithms (Table 6).

## USAGE NOTES

### Software Re-Use

For data preprocessing:

‐ FreeSurfer (http://surfer.nmr.mgh.harvard.edu/)
‐ FSL (https://fsl.fmrib.ox.ac.uk/fsl/fslwiki)
‐ MRTrix (http://www.brain.org.au/software/)
‐ GNU Octave (http://www.gnu.org/software/octave/)
‐ NIAK (Neuroimaging Analysis Kit; MATLAB toolbox) (https://www.nitrc.org/projects/niak/)

The software to run the simulations can be found here: https://github.com/the-virtual-brain

For data standardization:

‐ MNE-BIDS (https://mne.tools/mne-bids/dev/auto_examples/convert_eeg_to_bids.html)
‐ Bidscoin (https://github.com/Donders-Institute/bidscoin)
‐ BIDS validator (https://github.com/bids-standard/bids-validator)

## CODE AVAILABILITY

Image preprocessing and processing:

https://github.com/BrainModes/TVB-empirical-data-pipeline and https://search.kg.ebrains.eu/instances/Software/71265c9f-5fe3-40e3-a7e4-b2bb45b5ea6e for cloud computing.

We included all used code for simulations and BIDS conversions in the prepared data structure. We plan to publish this code after publication.

## ACKNOWLEDGEMENTS

We gratefully acknowledge the Gauss Centre for Supercomputing e.V. (www.gauss-centre.eu) for funding this project by providing computing time through the John von Neumann Institute for Computing (NIC) on the GCS Supercomputer JUWELS at Jülich Supercomputing Centre (JSC).

Additionally, computation of underlying data has also been performed on the HPC for Research cluster of the Berlin Institute of Health.

We thank Simon Rothmeier and Matthias Reinacher for helping with data collection and initial processing.

We acknowledge support by EU Horizon Europe program Horizon EBRAINS2.0 (101147319), Virtual Brain Twin (101137289), EBRAINS-PREP 101079717, AISN 101057655, EBRAIN-Health 101058516, EIC grant PHRASE 101058240, by the Digital Europe Programme TEF-Health (101100700), Shaiped (101195135), CoordinaTEF (101168074) German Research Foundation SFB 1436 (project ID 425899996); SFB 1315 (project ID 327654276); SFB 936 (project ID 178316478; SFB-TRR 295 (project ID 424778381); SPP Computational Connectomics RI 2073/6-1, RI 2073/10-2, RI 2073/9-1; DFG Clinical Research Group BECAUSE-Y 504745852, Berlin University Alliance OpenMake, the Virtual Research Environment at the Charité Berlin and EBRAINS Health Data Cloud and the Berlin Institute of Health and Foundation Charité.

JM acknowledges funding by the Deutsche Forschungsgemeinschaft (DFG, German Research Foundation) – Project-ID 424778381 – TRR 295.

## COMPETING INTERESTS

The other authors declare no competing interests.

## AUTHOR CONTRIBUTIONS

J.M.: writing the original draft of the manuscript, preparation and curation of the data, conversion of the data to BIDS standards and meta-data annotation with openMINDS, writing the manuscript. M.S.: writing parts of the original manuscript draft, data collection, data preprocessing, writing the manuscript, technical validation, development of the brain modeling software, proposition of BIDS specification for modeling data. P.R.: writing parts of the original manuscript draft, study setup, providing resources, data collection, data preprocessing, writing the manuscript, development of the brain modeling software. P.T.: simulations and fitting of the personalized brain network models, data analysis.

## Notes

### Summary of Updates

Funding acknowledgements updated and minor corrections

